# Encoding object-location memories along the proximodistal axis of CA1

**DOI:** 10.1101/2022.10.17.512601

**Authors:** Yusuke Teratani-Ota, Brian J. Wiltgen

**Author notes:** Correspondence: Brian J. Wiltgen.

## Abstract

The hippocampus is thought to combine “what” and “where” information from the cortex so that objects and events can be represented within the spatial context in which they occur. Surprisingly then, these distinct types of information remain partially segregated in the output region of the hippocampus, area CA1. In this region, objects preferentially activate neurons in the distal segment (adjacent to the subiculum) while spatial locations are precisely represented by neurons in the proximal segment (adjacent to CA2). This difference likely results from distinct anatomical connections; proximal CA1 receives direct input from the medial entorhinal cortex (which encodes spatial context) whereas distal CA1 has reciprocal connections with the lateral entorhinal cortex (which encodes objects and events). Based on these findings, it has been proposed that CA1 contains two distinct representations; one that encodes the animal’s spatial location and another that encodes objects that are present in the environment. The current study aimed to determine the role of distal CA1 in learning the location of objects in an environment. To do this, we first examined c-Fos expression in proximal and distal CA1 to see if we could replicate previous findings and confirm that neurons in these distinct segment are responsive to different stimuli. As previous studies indicate that catecholamines can regulate the activity of segments of CA1, we then investigate the role of catecholamines on learning object locations using 6-OHDA or SCH23390 to lesion catecholaminergic input and block D1/D5 receptors, respectively. Finally, we monitored calcium activity with fiber photometry while animals performed a hippocampal-dependent object location memory task.

## Introduction

The hippocampus (HPC) is thought to form episodic memories by integrating spatial and non-spatial inputs from the entorhinal cortex (EC). However, the way these distinct types of information are associated in different subregions of the HPC is not completely understood. Since the discovery of place cells, animal work has traditionally focused on the contribution the HPC makes to spatial learning and memory (McNaughton et al., 2006; O’Keefe, 1976). However, we now know that the HPC processes many different types of information. For example, time cells fire in precise temporal sequences, which is thought to allow spatial and non-spatial information to be linked across time (MacDonald et al., 2011).(Eichenbaum et al., 1999; Ekstrom & Ranganath, 2017). Recent studies have also demonstrated that CA1 pyramidal neurons respond to specific auditory cues, odors, and objects (Aronov et al., 2017; Burke et al., 2011; Igarashi et al., 2014; Nakamura et al., 2013). These responses reflect the diverse types of information the HPC receives from cortical and subcortical areas.

The entorhinal cortex (EC) is the major cortical input to the HPC. It can be subdivided into the medial entorhinal cortex (MEC) and the lateral entorhinal cortex (LEC), which are functionally distinct from one other. The MEC is important for forming allocentric representations of space via cells that fire in a grid-like pattern throughout the environment (Hafting et al., 2005; Hargreaves et al., 2005). In addition to grid cells, the MEC contains border cells, speed cells, and head-direction cells, all of which are important for spatial navigation (Giocomo et al., 2014; Kropff et al., 2015; Solstad et al., 2008). Cells in LEC are not spatially modulated but instead responsive to objects or odors in the environment. For instance, LEC neurons fire near objects that are encountered during an open foraging task (Deshmukh & Knierim, 2011). The removal of an object causes a different set of cells to fire at the site where it was previously located. This suggests the LEC stores and retrieves associations between objects and their spatial location (Tsao et al., 2013). Interestingly, LEC neurons show little to no firing in response to visual landmarks that make up the spatial environment (Yoganarasimha et al., 2011). Instead, this information is processed by the MEC and used to form allocentric representations of the environment that are necessary for spatial navigation.

The anatomical connections between the EC and the HPC have been characterized extensively. The MEC and LEC both send direct projections to granule cells in the dentate gyrus via the perforant path. Cells in DG transmit this information to pyramidal neurons in CA3 via the mossy fiber pathway. And finally, Schaffer collaterals from CA3 neurons terminate on cells in CA1. CA1 neurons also receive a direct project from the EC. This projection, called the temporoammonic pathway, bypasses DG and CA3 and synapses directly on CA1 neurons. Interestingly, the inputs from MEC and LEC are segregated along the proximodistal axis of CA1. This is not the case in DG and CA3, where individual neurons receive input from both regions.

The proximal segment of CA1 (adjacent to CA2) receives input from the MEC while the distal segment (adjacent to the subiculum) is targeted by neurons in the LEC (Masurkar et al., 2017; Witter et al., 2000). These segments then send return projections back to the deep layers of EC; proximal CA1 to MEC and distal CA1 to LEC. Based on this anatomy, it is hypothesized that the proximal CA1 determines the animal’s position in space, while distal CA1 encodes non-spatial stimuli in the environment, like objects and odors.

Consistent with this idea, *in vivo* electrophysiology recordings and IEG studies have shown that distal CA1 responds more strongly to the presence of objects than proximal CA1. In addition, the place cells in proximal CA1 (which respond to the animal’s location) are more precise and stable than those in distal CA1(Burke et al., 2011; Henriksen et al., 2010). We also showed that excitotoxic lesions of proximal, but not distal CA1, make it difficult for animals’ to recognize a familiar spatial environment (Nakazawa et al., 2016).

Pyramidal cells activity is strongly modulated by catecholamine released from locus coeruleus (LC) and ventral tegmental area (VTA) terminals in CA1. While the VTA is known to release dopamine in many brain areas, it was only recently discovered that the LC co-releases norepinephrine <u>and</u> dopamine in the HPC (Smith & Greene, 2012). In fact, catecholaminergic inputs to the HPC primarily come from LC neurons (Kempadoo et al., 2016; McNamara & Dupret, 2017; Wagatsuma et al., 2018). Most of the VTA projections to the HPC are GABAergic or glutamatergic and only 10% are dopaminergic. Although small in number, these inputs have been shown to stabilize hippocampal representations of space (McNamara et al., 2014). Catecholamine release in the HPC modulate entorhinal inputs in a bidirectional and frequency dependent manner. During low-frequency stimulation, dopamine attenuates the direct entorhinal input to CA1 (Ito, 2007, 2008). In contrary, under high-frequency stimulation protocols, dopamine potentiates entorhinal inputs in CA1. These effects have been observed in both proximal and distal CA1 (Ito et al., 2010). Unlike dopamine, norepinephrine primarily modulates entorhinal inputs to proximal CA1. This implies that dopamine and norepinephrine play distinct roles in filtering the transmission of sensory information from the entorhinal cortex to proximal and distal CA1.

Despite these data, it is still not known if activity in distal CA1 is required to learn and remember the position of objects in space. The majority of studies to date have simply recorded single unit activity in distal CA1while animals are exposed to objects. However, this activity has not been directly linked to memory performance (Burke et al., 2011; Deshmukh & Knierim, 2013; Hartzell et al., 2013; Ito & Schuman, 2012). In our study, we aimed to determine the role of distal CA1 in learning the location of objects in an environment. We started by examining c-Fos expression in proximal and distal CA1 to see if we could replicate previous findings and confirm that neurons in these distinct segment are responsive to different stimuli. As previous studies indicate that catecholamines can regulate the activity of segments of CA1, we then investigate the role of catecholamines on learning object locations using 6-OHDA or SCH23390 to lesion catecholaminergic input and block D1/D5 receptors, respectively. Finally, we monitored calcium activity with fiber photometry while animals performed a hippocampal-dependent object location memory task.

## Materials and methods

### Subjects

All procedures were approved by the Animal Care and Use Committee at UC Davis. For experiment 1, C57BL6 mice were used from in-house bred mice. All other experiments used F1 hybrids (C57BL/6N x 129S6; Taconic). Animals were all group-housed prior to all experiments. Mice were maintained on a 12/12 light/dark cycle and were given free access to food and water. All surgeries were performed at 8 weeks of age and behavioral experiments began at 10-12 weeks of age. For experiments not requiring surgeries, behavioral experiments began at 8-12 weeks of age.

### Viral construct

AAV9-CaMKII-GCaMP6f were packaged by the Vector Core at the University of North Carolina with a titer of 4.10×10^12^ viral particles/mL. The virus was diluted to 4.10×10^11^ with sterile PBS prior to surgical procedures.

### Stereotaxic surgeries

For experiment 2, mice were anesthetized with isoflurane (2-5%) and head-fixed into a stereotactic apparatus (Kopf instruments). A craniotomy was performed with a small precision drill and the virus was delivered with a glass pipette needle at an injection rate of 2nL per sec using a microinfusion pump (World Precision Instruments). 150μL of AAV9-CaMKII-GCaMP6f (Penn University) was infused in the HPC. To target proximal CA1, 150nL of virus was microinjected ([AP] –2.0 mm, [ML] ± 2.15mm, [DV] -1.2mm), followed by optic fiber implantation ([AP] –2.0 mm, [ML] ± 2.15mm, [DV] -1.0mm). To target distal CA1, 150nL of virus was microinjected ([AP] –2.0 mm, [ML] +0.75mm, [DV] -1.2mm), followed by optic fiber implantation ([AP] –2.0 mm, [ML] + 0.75mm, [DV] -1.25mm). For experiments 3 and 4, mice were bilaterally infused with 150μL of either sterile saline or 6-Hydroxydopamine (6-OHDA) hydrobromide (HelloBio, HB1889) in the HPC.

### Behavioral apparatus

An open-field box (37.5 × 37.5 × 37.5cm) made from white king starboard was used for the task. One of the walls was labeled with two vertical black stripes to serve as a spatial cue for the object-location memory task. An air purifier was turned on high to produce white noise in the room. Additionally, red lights were used for all behavioral experiments. For a novel context, a white PVC tube (37.5 × 37.5cm) was used with an air purifier was turned on low. Dim light settings were used to differentiate the context from the object-location memory task context.

### c-Fos experiments

Animals were all single-housed prior to the start of a behavioral experiment to prevent any factors that may induce activity unrelated to the task. Before the start of behavioral experiments, all animals were handled for 7 days (5-minutes each). The habituation sessions consisted of 15 minutes of exploration in an open-field environment for three consecutive days. For experiment 1, mice were randomly assigned to one of three conditions for the test session: control, object, and place. The control condition consisted of animals being placed in the habituated environment. For the object condition, animals were put into the habituated open-field box but with a pair of identical objects. Finally, animals assigned to the place condition were put into a completely novel environment (PVC tube). For experiment 3, the procedure was the same as experiment 1 except for the test session; the test session only consisted of the object condition.

### c-Fos experiments

Animals were all single-housed prior to the start of a behavioral experiment to prevent any factors that may induce activity unrelated to the task. Before the start of behavioral experiments, all animals were handled for seven days (5-minutes each). For experiment 1, C57BL6 mice at the age of 8-12 weeks of age were purchased from Jackson Laboratory. Mice were habituated to the open-field environment for three consecutive days (15-minutes each) and then randomly assigned to one of three conditions following habituation to an open-field box: re-exposure to context A (Control), exposure to novel objects in context A (Object), or exposure to a new context (Place). For experiment 3, C57BL6/129 F1 hybrid mice at the age of 6-8 weeks of age were purchased from Taconic Farms. Mice were randomly assigned to receive 6-OHDA or sham lesions. Following recovery, mice underwent the same behavioral procedure as experiment 1 except they only underwent the object condition. Animals in both experiments were transcardially perfused nintey minutes from the start of the testing session and fixed overnight in preparation for immunohistochemistry analysis. The amount of distance traveled for each session was measured using Noldus Ethovision tracking software.

### Immunohistochemistry for c-Fos

Ninety-minutes after testing, mice were transcardially perfused with 4% PFA. After a twenty-four hour post-fixation, 40uM coronal sections were collected and stained for c-Fos. Prior to all antibody staining, coronal slices were washed with 1X phosphate buffer solution (PBS) three times. Slices were incubated in donkey blocking buffer containing 2% donkey serum, 0.2% Triton X, and PBS for 15 minutes. Then, slices were incubated in c-Fos rabbit antibody for 24 hours at room temperature. A polyclonal rabbit antibody (1: 1000, Santa Cruz) was used for experiment 1 while a monoclonal rabbit antibody (1:5000, Cell signaling) was used for experiment 4. Following the primary antibody staining, slices were placed in biotinylated donkey anti-rabbit secondary antibody (1:500, Jackson ImmunoReseach) for sixty minutes at room temperature, followed by streptavidin-Cy3 (1:500, Jackson ImmunoResearch) for forty-five minutes). Finally, slices were DAPI stained for fifteen minutes and mounted on slides.

### Novel object location memory tasks

For these experiments, C57/129 F1 hybrid mice were purchased from Taconic Farm. Mice were placed into an open-field box for three-consecutive days for two-minutes each session. One of the walls was labeled with two vertical black stripes to serve as a spatial cue. During the sample session, mice were placed into the open-field box with a pair of identical objects located in front of the spatial cue for fifteen minutes. After a twenty-four hour delay, one of the two objects was moved to the opposite wall to test the animal’s memory for ten-minutes. The placement of objects was counterbalanced. The discrimination index was calculated by dividing the exploration for the novel object by the total exploration of objects (stationary and displaced). A discrimination index of 0.5 will indicate performance at chance level. Due to the nature of the task, only the first five minutes was used to calculate the discrimination index.

### SCH23390 behavioral experiments

For all experiments, SCH23390 (Tocris Bioscience) was used to block D1/D5 receptors. The drug was prepared by dissolving 10mg of SCH23390 in 10mL of sterile saline to prepare a 1mg/mL concentration. Mice were administered SC23390 by IP injection or intracranial infusion. For IP injections, mice received the drug thirty minutes prior to the sample session and allowed to freely move in their home cage. For intracranial infusions, 0.5μL of SCH23390 were administered via bilateral infusions (0.1μL/min) using a 10μL Hamilton syringe fifteen minutes before the sample session. Animals freely explored an empty cage during infusion of the drug. Following infusion, the injectors were left in place for two additional minutes before removing it from the cannulae. Animals remained in the empty cage until the sample session.

The discrimination index was calculated by dividing the exploration for the novel object by the total exploration of objects (stationary and displaced). A discrimination index of 0.5 will indicate performance at chance level. Due to the nature of the task, only the first five minutes was used to calculate the discrimination index.

### Fiber photometry and analysis

Following recovery from surgical procedures, animals were handled for seven-days by attaching the patch cord to the optic fiber implant. Animals were allowed to freely walk around an empty cage for five minutes each day. After handling, animals performed the OLM task as described above while monitoring calcium activity using a basic fiber photometry setup (Doric). To monitor the isosbestic and GCaMP6f signals, wavelengths of 405nm and 465nm was used. Before animals were placed in the open-field box, a ten-minute recording session took place to allow photobleaching of signals. The Noldus Ethovison software was connected to Doric’s system to send TTL pulses whenever an animal explored the objects. The TTL pulses were used to identify the time points in which the animals explored different types of objects. Analysis of all photometry data was performed using the open-source Guided Photometry Analysis in Python (GuPPY) to create peri-stimulus/event time histograms (PSTH) and calculate the area under the curve (AUC) during object exploration (https://github.com/LernerLab/GuPPy).

### Immunohistochemistry for TH

Separate 40um coronal slices from the same subjects from experiment 3 were collected to stain for tyrosine hydroxylase (TH). Prior to all antibody staining, coronal slices were washed with 1X phosphate buffer solution (PBS) three times. Slices were incubated in donkey blocking buffer. Next, slices were incubated in TH polyclonal rabbit antibody for 24 hours at room temperature. The following steps after the incubation of primary antibody is the same procedure as described in the c-Fos immunohistochemistry protocol.

### Imaging and quantification

Samples were imaged using an Olympus fluorescence virtual slide scanning microscope. For experiment 1, z-stack images were obtained for 40um slices. Fluorescent images were then imported into FIJI as a grayscale image. Each image’s channel was separated to hand count c-Fos positive cells through z-stacks. A 3D Objects Counter macro in FIJI was used to estimate the number of DAPI stained nuclei in each area by dividing the obtained volume by the average single nucleus volume for the area. To obtain the percentage of c-Fos positive cells, the number of c-Fos positive cells were divided by the total number of cells. For all other experiments, 40um sections were imaged using a single plane scan. To quantify c-Fos, CA1 segments were cropped to an area of 30,000um^2^ and imported to a grayscale image into FIJI. A cell count macro developed by Kyle Puhger was then used to process the image (https://github.com/kpuhger/imagej_cellcount). Briefly, the macro enhanced contrast of the image, removed outlier pixels (set by the user), then a gaussian blur is applied, and finally the maxima is identified. To quantify TH-expression, segments were cropped to an area of 30,000um^2^. After importing images to FIJI, images were cropped to an area of 30,000um^2^ for each segment of CA1. Images were then processed by using despeckle, removing outliers, and subtracting background. The mean intensity of the image was then calculated to quantify TH-expression.

### Statistics

c-Fos expression differences were analyzed with a two-way repeated measures ANOVA and post-hoc comparisons were made using Tukey’s test. Activity levels during sample session and discrimination index were analyzed with a two-tailed unpaired t-test. For fiber photometry, the AUC observed in the sample session was analyzed using a two-tailed unpaired t-test. For the test session, a two-way repeated measures ANOVA and a Tukey’s test was used for post-hoc comparisons.

## Results

### Experiment 1: Examining what kinds of information are processed along the proximodistal axis of CA1

Previous studies found proximal and distal CA1 responsive to spatial and non-spatial stimuli, respectively. To replicate previous findings, we habituated animals to an open-field environment for three consecutive days. Following habituation, animals were either placed in a habituated environment (control), exposed to a pair of identical objects in the habituated environment (object), or placed in a novel environment (place) (Figure 1A). After the test session, we examined c-Fos as a proxy for neural activity along the proximodistal axis of CA1 to determine what types of stimuli each segment processes.

**Figure 1.**
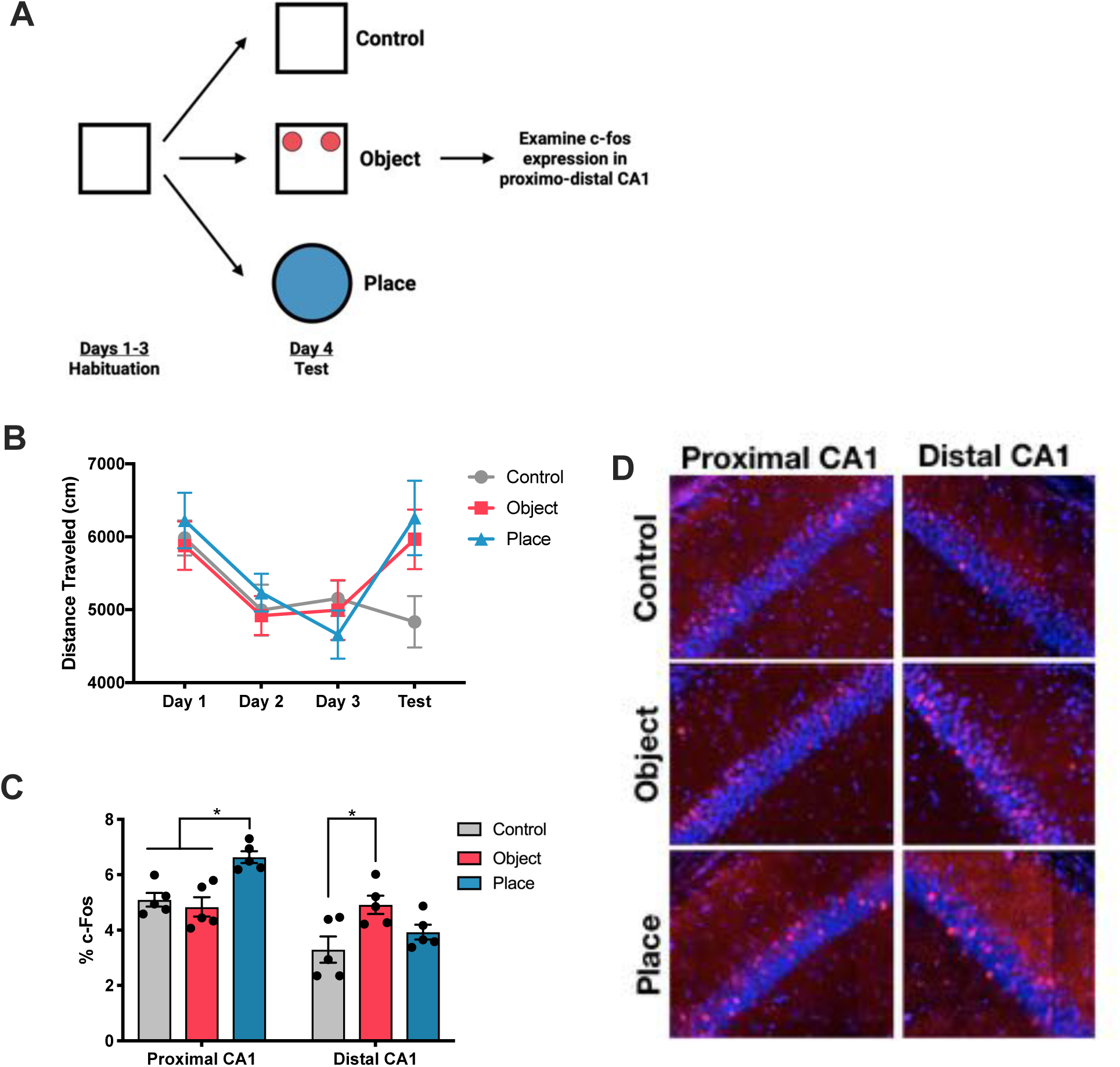
Proximal and distal CA1 are modulated by different stimuli. **(A)** Experimental design. Animals were habituated to an open-field environment for fifteen minutes per day for three days. Following habituation, animals were placed in the habituated open-field environment (control), exposed to novel objects (object), or put into a novel place (place). **(B)** All groups habituated to the environment across habituation. Activity levels increased when mice were exposed to novel objects or a new place (2-way repeated measures ANOVA, Interaction F(6, 36) = 3.707, p >0.05). **(C)** Quantification of c-Fos expression in proximal and distal CA1. Mice exposed to a new place displayed higher levels of c-Fos in proximal CA1 while mice exposed to novel objects had higher levels of c-Fos in distal CA1 (2-way repeated ANOVA, Interaction F(2, 12) = 12.18, p <0.01). **(D)** Representative images of c-Fos expression in proximal and distal CA1 from mice in each condition (n = 5 per group)

As expected, animals in all groups showed a gradual decrease in the amount of distance traveled in the environment during habituation (Figure 1B; Days 1-3). Although there was a significant interaction of days and condition, post-hoc tests did not reveal any significant differences (2-way repeated measures ANOVA, Interaction F(6, 36) = 3.707, p >0.05). However, it is still worthwhile to observe that animals exposed to objects or a novel place on the test day displayed increased activity levels. Quantification of c-Fos expression revealed elevated activity in proximal CA1 of mice that were exposed to a novel place compared to control mice and those exposed to objects. However, animals exposed to objects or a novel place displayed higher activity levels relative to the control condition. We also observed elevated c-fos activity in proximal CA1 of mice exposed to a novel place compared to mice exposed to the control or object conditions (Figures 1B & 1C). In contrast, mice that were exposed to novel objects had higher c-Fos activity in distal CA1 compared to that of the place and control conditions (2-way repeated measures ANOVA, Interaction F(2, 12) = 12.18, p <0.01, Tukey’s multiple comparison p < 0.01). Despite animals in the object and place conditions displaying similar activity levels during testing, we observed differential activity patterns along the proximo-distal axis of CA1.

Thus, our observation cannot simply be due to animals being more active in their assigned conditions. These results replicate previous studies suggesting that proximal CA1 is more spatially tuned while distal CA1 is responsive to objects.

### Experiment 2: Catecholaminergic inputs role in regulating proximodistal CA1 during object related memory tasks

Catecholamines have been shown to be important in regulating entorhinal inputs along the transverse axis of CA1 (Ito, 2007; Ito et al., 2010; Ito & Schuman, 2012). We next investigated whether catecholamines are responsible for the object induced c-Fos expression we observed in our previous experiment. To do this, a neurotoxin 6-hydroxydopamine (6-OHDA) hydrobromide was bilaterally infused into the HPC to lesion all catecholaminergic inputs. After recovery, animals were placed in an open-field environment for three consecutive days and then exposed to a pair of identical objects during the testing session (Figure 2A). We then performed immunohistochemistry to examine c-Fos activity in proximodistal CA1 following the behavioral procedure. To confirm 6-OHDA lesions, we quantified tyrosine hydroxylase (TH) expression in proximal and distal CA1 via immunohistochemistry (Figure 2B and 2C). Animals with 6-OHDA lesions had significantly lower TH-positive axons in the HPC in both segments of CA1 relative to the vehicle condition (2-way repeated measures ANOVA, Interaction F(1, 9) = 24.60, p < 0.01). The slight reduction of TH expression observed in proximal CA1 may be driven due to high background from immunostaining; comparison of TH-expression images from each condition shows a clear difference in proximal and distal CA1 (Figure 2C). Analysis of c-Fos expression revealed that there were main effects of lesion conditions and CA1 segments (2-way repeated measures ANOVA, main effect of lesion condition: F(1, 10) = 8.812, p < 0.05; main effect of segment: F(1, 10) = 90.12, p < 0.05) but there were no significant interactions observed (2-way repeated measures ANOVA, Interaction: F(1,10) = 0.6095, p > 0.05). This suggests that catecholamines may be important for increasing c-Fos expression driven by object exploration but does not differentially modulate activity along the proximodistal axis of CA1.

**Figure 2.**
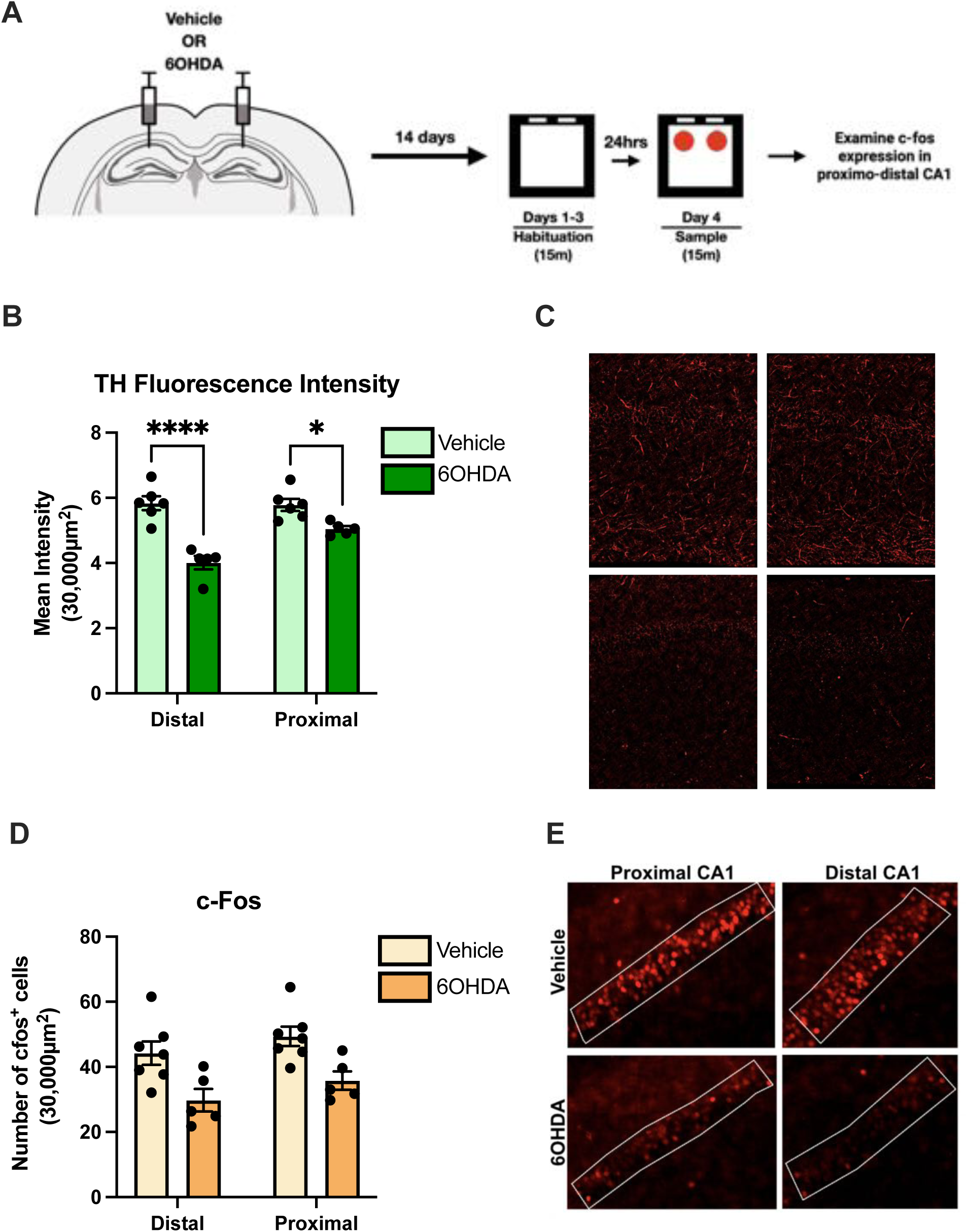
6-OHDA catecholaminergic lesions reduced c-Fos expression in proximal and distal CA1. **(A)** Experimental design. Animals received bilateral infusions of 6-OHDA or saline in the HPC. After recovery, animals habituated to an open-field environment for three consecutive days (fifteen min per day). The following day, both groups of animals were exposed to novel objects. Ninety-minutes after, animals were sacrificed, and brain tissue was fixed to immunostained for c-Fos. **(B)** Quantification of tyrosine-hydroxylase (TH) in HPC. A reduction of TH-positive axons in proximal and distal CA1 were observed in the 6-OHDA group (2-way repeated measures ANOVA, Interaction F(1, 9) = 24.60, p < 0.01). Only a slight reduction was observed in proximal CA1, but this may be due to high background in images (See figure 4C) **(C)** Representative images of TH expression in proximal and distal CA1. **(D)** Quantification of c-Fos expression in proximal and distal CA1. Catecholaminergic lesions with 6-OHDA reduced overall c-Fos expression in proximal and distal CA1 (2-way repeated measures ANOVA, main effect of lesion condition: F(1, 10) = 8.812, p < 0.05; main effect of segment: F(1, 10) = 90.12, p < 0.05). No interaction between the CA1 segment and the group condition were found (2-way repeated measures ANOVA, Interaction: F(1,10) = 0.6095, p > 0.05). **(E)** Representative images of c-Fos expression in proximal and distal CA1 of 6-OHDA and vehicle animals.

To examine the contribution of catecholamines on object-related memories, mice with catecholaminergic lesions performed the hippocampal-dependent object location memory (OLM) task. In this task, animals habituated to an open-field environment for three consecutive days. The following day, animals were presented with a pair of identical objects in the habituated environment. After a 3-hour delay, one of the presented objects was moved to a new location and animals were given time to explore (Figure 3A). As mice are naturally inclined to explore novelty, they should spend more time exploring the object that moved to a new location. Although LC inputs in the HPC have been shown to be important for learning new contexts (Wagatsuma et al., 2018), we found no differences in activity levels during habituation to the open-field environment (2-way repeated measures ANOVA, Interaction F(2, 46) = 1.198, p > 0.05) (Figure 3B). At testing, mice with 6OHDA lesions had impaired discrimination when compared with the vehicle group (t(23) = 2.958, p < 0.05) (Figure 3D). The deficit cannot be explained by differences in exploration behavior during the sample session as both groups explored objects in a similar manner (t(23) =0.2705, p >0.05) (Figure 3C).

**Figure 3.**
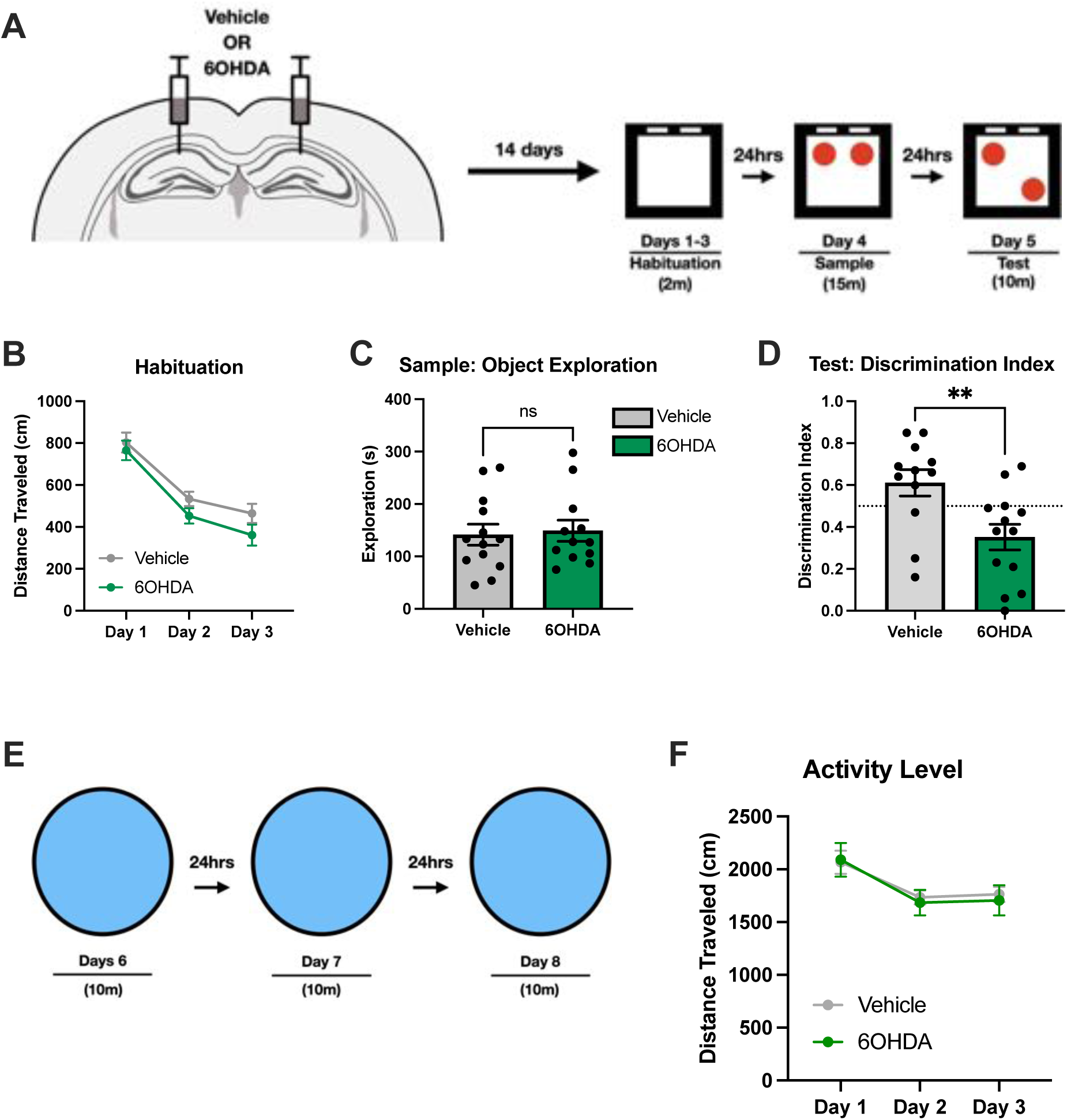
Catecholaminergic inputs are required for appropriate object discrimination. **(A)** Experimental Design of OLM. 6-OHDA or saline was bilaterally infused in the HPC and performed the OLM task following recovery. **(B)** Both groups habituated to the environment across the days. Activity levels between both groups were similar throughout habituation (2-way repeated measures ANOVA, Interaction F(2, 46) = 1.198, p > 0.05). **(C)** Total object exploration during sample session. No differences were observed between the 6-OHDA and vehicle group (t(23) =0.2705, p >0.05). **(D)** Performance of OLM measured by discrimination index at testing. Animals with 6-OHDA lesions had impaired discrimination relative to the vehicle group (t(23) = 2.958, p < 0.05). **(E)** Experimental design of habituation task. The same animals were exposed to a novel place for three consecutive days, fifteen-minutes per day. **(F)** Both groups habituated to the novel place in a similar manner (2-way repeated ANOVA, Interaction F(2,46) = 0.4278, p >0.05).

Deficits may not have been observed during the habituation sessions from the OLM task as animals were only given a short time frame (2 minutes) to explore the environment. Differences may have been observed if animals are given a longer period to explore. The same animals from the OLM experiment performed a habituation task in a novel environment; animals were allowed 10 minutes to explore a novel environment for three consecutive days (Figure 3G). No differences were observed in the activity levels across days of habituation in both groups, suggesting that catecholaminergic inputs are not required for learning new contexts (2-way repeated ANOVA, F(2,46) = 0.4278, p >0.05) (Figure 3F). Due to 6OHDA lesions being a chronic manipulation, there is sufficient time for changes in the brain to occur to compensate for the lack of catecholaminergic inputs in the HPC. However, this compensation is not sufficient to allow animals to properly learn object-related information for the OLM task. As 6-OHDA lesions prevent the release of both dopamine and epinephrine, we cannot make any conclusions about which catecholamine is important for processing object-related information.

### Experiment 3: Determining whether dopamine is important for object-related learning

The D1/D5 dopamine receptors are widely expressed in the HPC and have been demonstrated to play a crucial role in the acquisition of new information by regulating LTP/LTD (Lemon & Manahan-Vaughan, 2006; Otmakhova & Lisman, 1996, 1998). To determine if the blockage of D1/D5 receptors could explain the deficit we observed in our 6-OHDA experiment, we administered SCH23390 hydrochloride via IP injections prior to the sample session in the OLM task (Figure 4A). Administration of SCH23390 resulted in hypoactivity during the sample session (t(25) =2.958, p < 0.05) (Figure 4B). Due to a reduction in activity levels, mice were not able to freely explore the objects presented to them. This may not be surprising as the drug was administered systemically, which could affect motor systems that heavily rely on dopamine for initiating movement.

**Figure 4.**
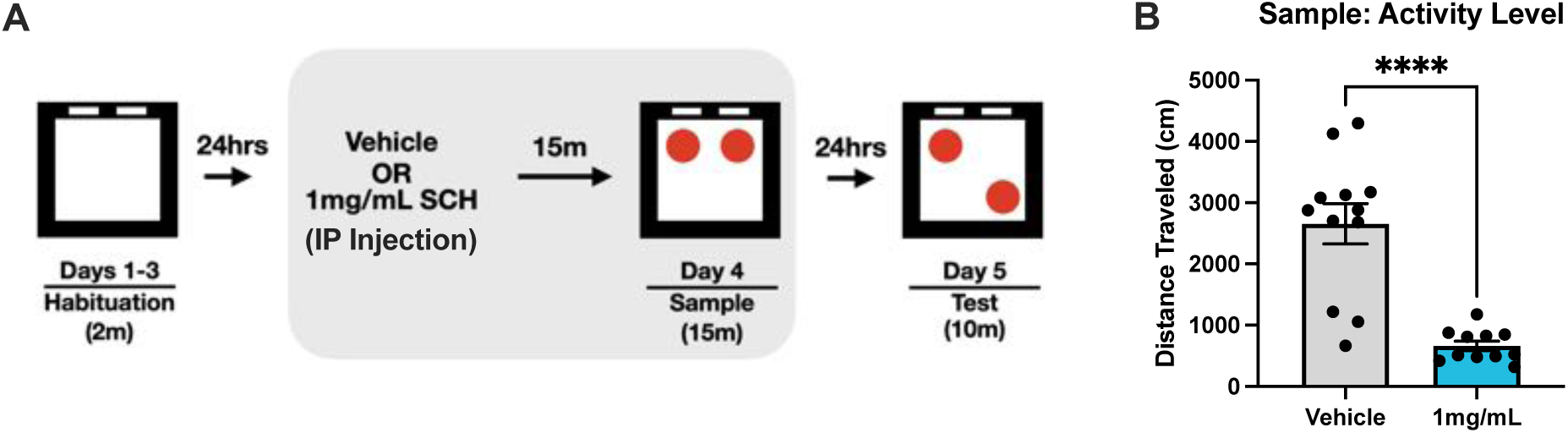
Administration of D1/D5 antagonist SCH23390 induces hypoactivity. **(A)** Experimental design. Animals performed the OLM task using the same procedure described in other experiment, except SCH23390 (1mg/mL) was administered via IP injection prior to the sample session. **(B)** Administration of SCH23390 induced hypoactivity during sample session (t(25) = 2.958, p< 0.05; t(25) = 2.413, p <0.05). Many animals were stationary after placing them in the open-field environment.

To circumvent the off-target effects of SCH23390, we bilaterally implanted cannulae above the CA1 region of the HPC. Administering SCH23390 directly to the HPC should prevent any motor deficits we observed during systemic injection of the drug. Although intracranial infusions did not reduce motor activity to the same extent of IP injections, animals administered with SCH23390 still displayed a significant decrease in overall activity levels and exploration behavior relative to the vehicle group (t(25) = 2.958, p< 0.05; t(25) = 2.413, p <0.05, Figure 6B & 6C). In addition, both groups of mice performed the OLM task around chance level at testing (t(25) = 0.7396, p > 0.05). The procedure for bilateral cannulae implants could have affected the vehicle group’s performance in the OLM task. Therefore, a different OLM protocol may have been necessary to allow animals to show appropriate levels of discrimination. However, regardless of the animal’s performance in the test session, it is still a concern that the drug affects the overall activity levels of the animals. Due to the side effects of SCH23390, it is difficult to determine whether dopamine is important for object-related memories. Future studies will examine the effects of a beta-adrenergic blocker, propranolol, to investigate whether norepinephrine is important for object-related processing.

**Figure 5.**
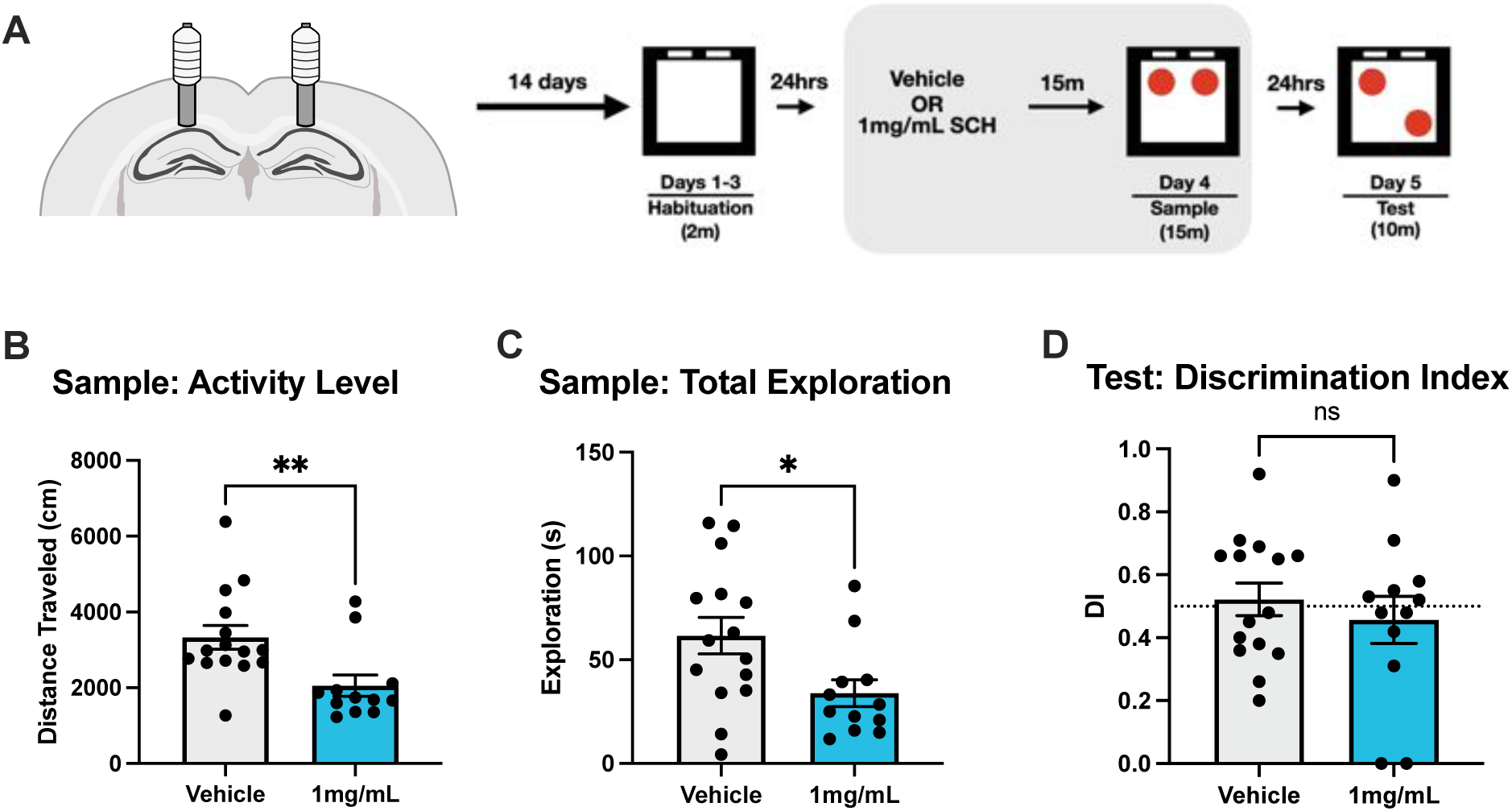
Intracranial infusion of SCH23390 reduces activity and object exploration behavior. **(A)** Experimental design. Cannulae were bilaterally implanted above CA1 to intracranially infuse 500μL of SCH23390 (1mg/mL) prior to the sample session of OLM. This method of drug administration was used to circumvent hypoactivity observed in the previous experiment. **(B)** Animals with intracranial SCH23390 infusion exhibited hypoactivity during the sample session (t(25) = 2.958, p< 0.05; t(25) = 2.413, p <0.05). The reduction in activity levels was not to the same degree of animals that received the drug via IP injections. **(C)** Performance at testing was impaired in both groups of animals (t(25) = 0.7396, p > 0.05). Bilateral cannulae implants may have disrupted animal’s normal processes due to the surgical procedure. Regardless of the results, it is a concern that SCH23390 affects overall activity levels of animals.

**Figure 6.**
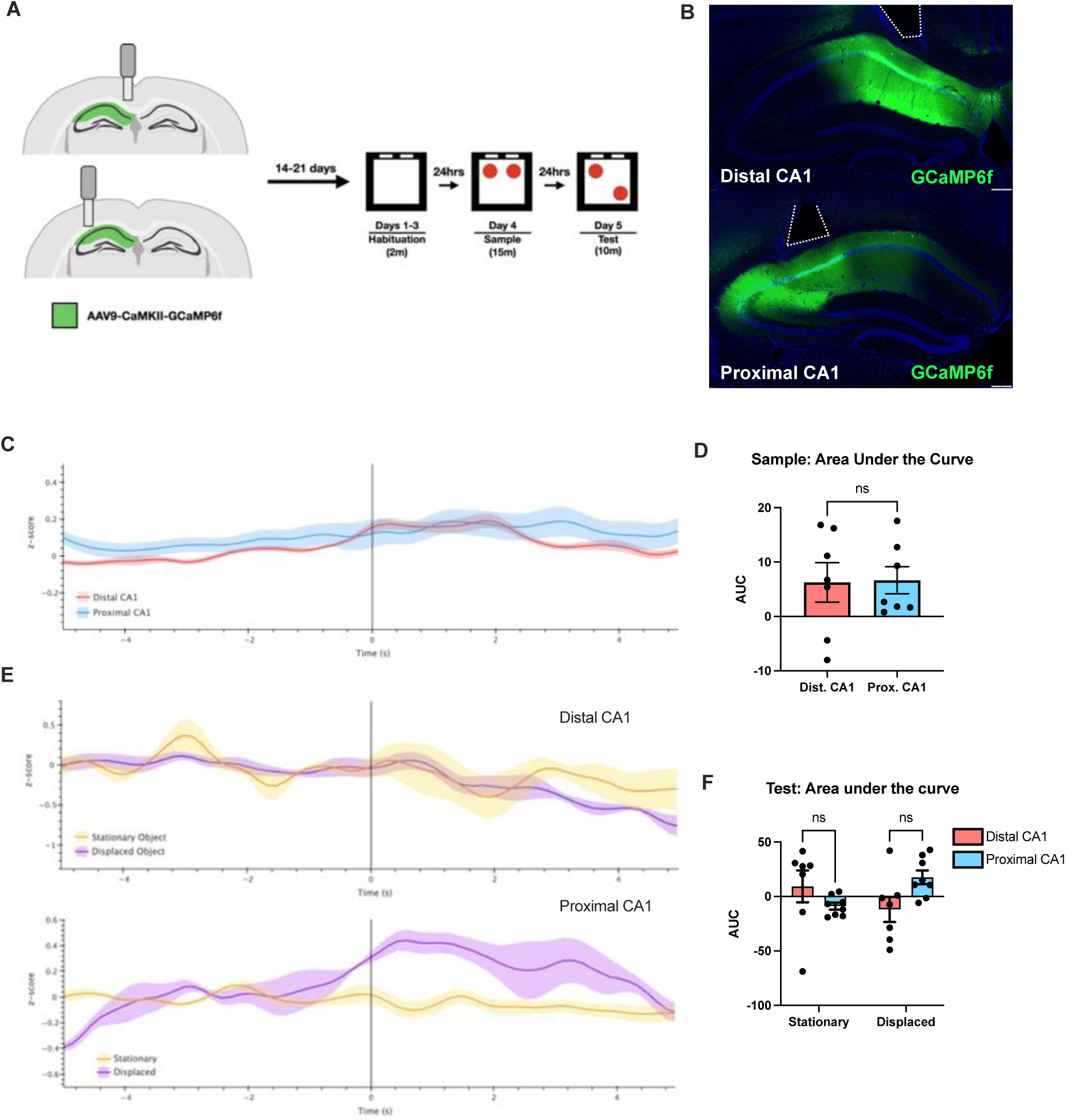
Proximal and distal CA1 have similar calcium activity during object exploration. **(A)** Experimental design. AAV9-CaMKII-GCaMP6f was unilaterally infused in the hippocampus and fiber optics were implanted above proximal or distal CA1. Animals performed the OLM task to monitor calcium activity during the sample and test sessions. **(B)** Representative image of viral expression of AAV9-CaMKII-GCaMP6f. **(C)** Peri-stimulus/event time histograms (PSTH) during sample session. Proximal (blue line) and distal (red line) CA1 had similar levels of activity during object exploration. **(D)** Quantification of area under the curve (AUC) during object exploration. No differences were observed between proximal and distal CA1 during exploration of objects (t(12) = 0.09, p >0.05). **(E)** PSTH of proximal (top) and distal (bottom) CA1 during exploration of the stationary (yellow line) and displaced object (purple line). **(F)** Quantification of AUC during exploration of stationary and displaced objects. There was a significant interaction between the type of objects and segments of CA1 but post-hoc analysis did not reveal any differences *2-way repeated ANOVA, Interaction F(1,13) =14.11, p<0.05). Calcium activity was higher when animals explored the stationary and displaced objects for distal and proximal CA1, respectively.

**Figure 7.**
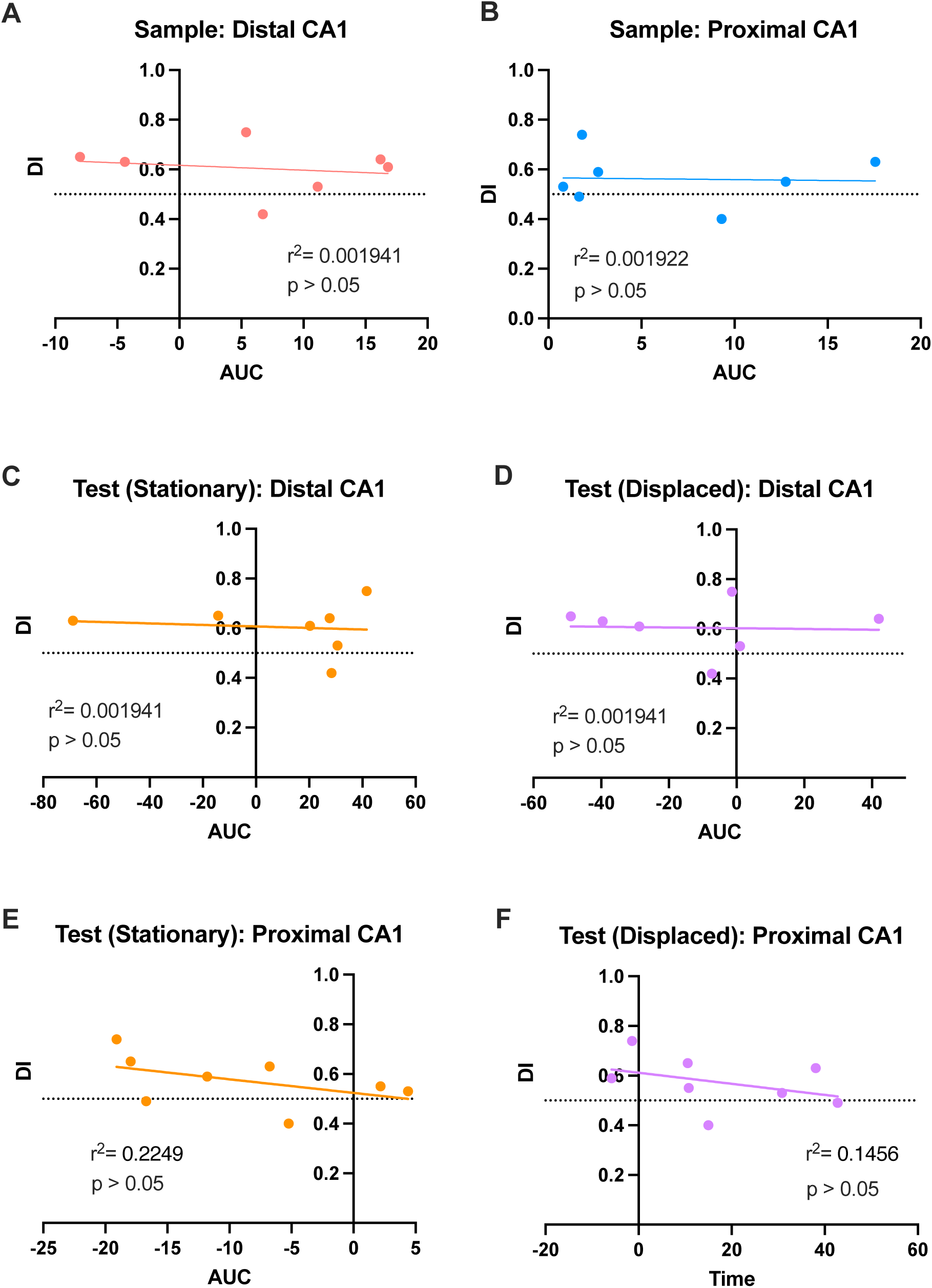
Calcium activity of proximal and distal CA1 do not correlate with animal’s performance on the OLM task. No significant correlations were found between the performance of the OLM task and the activity quantified by AUC during object exploration **(A)** Correlation of distal CA1 activity during the sample session with the discrimination index (r(5) = -0.18, p > 0.05). **(B)** Correlation proximal CA1 activity during the sample session with the discrimination index (r(5) = -0.04, p > 0.05). **(C)** Correlation of distal CA1 activity during exploration of stationary object with the discrimination index (r(5) = -0.11, p >0.05). **(D)** Correlation of distal CA1 activity during exploration of displaced object with the discrimination index (r(5) = -0.47, p > 0.05). **(E)** Correlation of proximal CA1 activity during exploration of stationary object with the discrimination index (r(5) = -0.04, p > 0.05). **(F)** Correlation of proximal CA1 activity during exploration of displaced object with the discrimination index (r(5) = -0.38, p>0.05).

### Experiment 4: Monitoring calcium activity of proximodistal CA1 during the object location memory task

Distal CA1 neurons display selective activity when animals explore different objects in an environment; however, the activity of these neurons fire to all objects regardless of its identity. It is unclear if distal CA1 exhibit distinct activity during an object-related memory task. To do so, we unilaterally injected an AAV9-CaMKII-GCaMP6f into CA1 and implanted fiber optics above proximal or distal CA1. We then monitored calcium activity while animals performed the OLM task.

To examine differences in calcium activity between proximal and distal CA1 during object exploration, we analyzed the area under the curve (AUC) during exploration. During the sample session, the activity of proximal and distal CA1 did not differ from each other when animals explored the objects (t(12) = 0.09, p >0.05). We next examined whether exploration of stationary or displaced objects resulted in different patterns of activity in segments of CA1. There was a significant interaction between the type of objects explored and the distinct segments of CA1 (2-way repeated ANOVA, Interaction F(1,13) =14.11, p<0.05). Although post-hoc comparisons were not significant, we found calcium activity in distal CA1 to be higher when animals explored the stationary object compared to the displaced object. In the contrary, calcium activity in proximal CA1 was larger when animals explored the displaced object (Figure 2F). These findings may indicate that proximal and distal CA1 process different aspects of information regarding the location of objects in an environment.

As there is variability in the animal’s performance of the OLM task, it may be possible that the performance is related to the activity of distal CA1 observed during the sample or testing sessions. The activity of either segment of CA1 during the sample session did not correlate to the performance of the OLM task (Distal CA1: r(5) = -0.18, p >0.05 and Proximal CA1: r(5) = -0.04, p>0.05). For the testing session, activity of proximodistal CA1 when animals explored the stationary object was not correlated to the discrimination index (Distal CA1: r(5) = -0.11, p >0.05 and Proximal CA1: r(5) = -0.47, p>0.05). The discrimination index and the activity of proximal and distal CA1 during exploration of the displaced object also were not correlated with each other (Distal CA1: r(5) = -0.04, p >0.05 and Proximal CA1: r(5) = -0.38, p>0.05). Overall, the performance of each animal could not be explained by differences in calcium activity monitored during object exploration (Figure 3). Fiber photometry monitors bulk calcium activity in the region of interest and as a result, it may be difficult to observe behaviorally relevant responses due to the amount of noise in the HPC. The HPC receives many inputs from cortical regions and thus, a technique that allows the recording of single neurons may be beneficial to observing activity of proximodistal CA1 during a learning task.

## Discussion

Here, we observed differential activity patterns along the proximodistal axis of CA1. Proximal CA1 showed an increase in c-Fos activity after exposure to a novel place. In the contrary, animals exposed to objects had elevated levels of c-Fos in distal CA1. This result is consistent with previous findings suggesting that proximal CA1 is spatially tuned while distal CA1 is responsive to objects. Despite the overall activity between the object and place conditions being similar, we observed distinct c-Fos expression at each segment of CA1. Thus, the pattern we observed is driven by the type of stimuli instead of increased activity levels.

We also found that catecholaminergic lesions in the HPC greatly reduced c-fos activity in proximal and distal CA1 following exposure to objects. Furthermore, animals without catecholaminergic inputs had impaired discrimination during the testing of the OLM task. Our results are consistent with previous studies suggesting that catecholamines are important for object-related memory tasks (Kempadoo et al., 2016; Moreno-Castilla et al., 2017). Surprisingly, our manipulation did not affect animals’ ability to learn about new contexts. Wagatsuma et al. had previously demonstrated that LC projections to CA3, but not CA1 or DG, were necessary for acquisition of a contextual memory (Wagatsuma et al., 2018). In our study, catecholaminergic lesions were mostly targeted to the CA1 subregion of the hippocampus and thus we may not have observed a deficit in the animal’s contextual memory. Catecholamines in the CA1 may not be necessary for the encoding of new places; however, release of catecholamines can result in the enhancement of memories by modulating consolidation processes (Duszkiewicz et al., 2019; Kempadoo et al., 2016; Takeuchi et al., 2016; Tsetsenis et al., 2021).

As 6-OHDA lesions eliminates all catecholaminergic inputs, we aimed to determine which catecholamine is responsible for OLM deficits we observed in our study. Blockage of D1/D5 receptors via administration of SCH23390 prior to the sample session resulted in hypoactivity of mice and thus reduced exploration behavior. By anecdotal observation, we did not observe any motor impairments in mice that was administered the drug. Animals given SCH23390 moved in a similar manner to the vehicle group when placed in the open-field environment. After a couple of seconds of exposure, animals with SCH23390 no longer explored the environment and only moved its head. The drug may not have affected the animal’s motor function but instead led to a decrease in their motivation to explore novel objects in the environment.

From our study, no conclusions can be made about whether dopamine is important for object-related memories from our studies. However, some studies provide evidence that dopamine may be the key neuromodulator for allowing animals to learn the OLM task. Kempadoo et al. (2016) optogenetically stimulated the LC in mice that received an intracranial infusion of SCH23390 or propranolol, a beta-adrenergic receptor antagonist, prior to the sample session of a weak training protocol. Under the weak training protocol, animals had poor discrimination, but this was mitigated by optogenetically stimulating the LC. The enhancement in performance was blocked with SCH23390 but not propranolol, implicating that norepinephrine may not be necessary for learning about the location of objects. Although norepinephrine may not be involved in the acquisition of an object memory, *in vivo* microdialysis of the HPC demonstrated that there is a 2-to-3-fold increase in dopamine and norepinephrine release when an object was displaced during testing of the OLM task. Therefore, norepinephrine may have a crucial role in the retrieval of an object-related memory or in detecting changes in the environment. The involvement of LC under normal learning conditions is still an open question. As catecholamines regulate the information flow of entorhinal inputs in CA1, it is important to understand how LC supports learning about an object’s location in the environment. The activity of LC is more homogenous relative to the HPC, and thus, fiber photometry may be sufficient to monitor dynamic changes that can occur with the animal’s exploration behavior.

Our fiber photometry experiments did not corroborate with previous findings regarding proximodistal CA1 having differential responses during object exploration. Calcium activity in each segment also did not correlate with the performance of each animal. In our c-fos study examining what types of stimuli proximodistal CA1 are sensitive to, only 4-5% of cells were found to be positive for c-Fos. The percentage of c-Fos positive cells does not necessarily equate to the absolute number of active cells. Many cells that do not express c-Fos can still fire action potentials (Tanaka et al., 2018). As fiber photometry monitors bulk calcium activity, it is possible that behaviorally relevant calcium signals cannot be observed due to other unrelated activity occurring in the background (Gunaydin et al., 2014). Due to the diverse information the HPC processes, fiber photometry may not have been the appropriate tool to examine dynamic changes that occur in the OLM task. Differences in proximal or distal CA1 could have been observed if techniques such as single-cell recording or miniscope calcium imaging were used to determine how individual cells may respond to object-related information.

It is also possible that to observe changes at the population level, animals may need to be engaged in a task that requires the association of a non-spatial stimuli with an outcome. The OLM task purely relies on the animal’s innate motivation to explore novelty. In an ethological setting, if a novel stimulus does not provide any valuable information, animals may be less inclined to learn anything about it. Using *in vivo* electrophysiology, Igarashi et al. (2014) examined the development of odor-specific representations in distal CA1 during a task that required mice to learn an association of a reward with a specific odor. During sampling of an odor cue, an oscillatory activity in the local field potential of distal CA1, but not proximal CA1, was observed at 20-40Hz. Furthermore, analysis of individual spikes in distal CA1 demonstrated that many neurons exhibited selective activity during the exposure to the reward-predicting odor as the mice successfully learned the association. Thus, it may be critical for a non-spatial stimulus to be associated with a relevant outcome to observe significant increases in bulk calcium activity in distal CA1.

Lastly, no differences could have been observed in our photometry experiment because both segments of CA1 can fire in response to objects; each segment may process different aspects of object exploration. As stated before, proximal and distal CA1 are typically thought to process spatial and non-spatial information, but this may be an oversimplification of the role of these two circuits within the HPC. While grid cells in MEC maintain their grid-like firing properties regardless of whether objects are present, it has been reported that there is a different type of cells called an object-vector cell that fires when an animal is facing an object at a specific orientation and a certain distance away from an object (Høydal et al., 2019). The firing properties of object vector cells remain the same even if the environment, the object, or the object’s location is different. Therefore, it is a purely allocentric representation of the animal’s position relative to an object in any environment. This representation allows the animal to navigate using landmarks to reach a goal destination. Object-vector cells have yet to be found in proximal CA1 but as it mostly receives projections from the MEC, there are likely cells that show stable firing when an animal is at a certain distance from an object.

A recent study observed place cell activity in proximodistal CA1 when animals explored objects during the OLM task (Vandrey et al., 2021). In addition to place cells, trace cells or cells that fire to an object’s previous location were also found in proximodistal CA1. Although object-related activity was found in both segments, cells in distal CA1, but not proximal, exhibited higher spatial specificity and were strongly modulated by the proximity of objects. For instance, trace cell activity of distal CA1 fired precisely to the displaced object’s previous location than proximal CA1. Instead of firing precisely to the object’s previous location, trace cell activity in proximal CA1 may be influenced by the animal’s head-direction and the animal’s distance from the object like object vector cells. These studies may explain why exposure to objects elicited similar levels of c-Fos expression in proximal and distal CA1. Proximal CA1 processes the precise location of an animal in relation to the object’s location, while distal CA1 encodes the location of objects in an environment, which is consistent with Knierim’s new framework of the parallel circuits within the hippocampal system (Knierim et al., 2014). The new framework describes the MEC-proximal CA1 circuit as being imperative for storing a spatial framework of a context, allowing animals to determine where they are in the environment. Animals may use distal cues or objects/landmarks to navigate to a goal of interest. In contrast, the LEC-distal CA1 circuit is important for storing information about the content of an experience. This can consist of individual items in the environment, what certain objects are, and the location of objects in the environment. Due to the diverse responses in the HPC, future studies will use techniques such as miniscope calcium imaging to record activity of individual neurons during the OLM task to better understand how proximodistal CA1 is involved in object-related memories. In addition, it will be of interest to utilize catecholaminergic biosensors (such as Dlight, GRAB_DA_, or GRAB_NE_) with calcium sensors to examine how dopamine or norepinephrine influence the activity of cells that respond to objects in each segment.

